# dScaff – an automatic bioinformatics framework for scaffolding draft *de novo* assemblies based on reference genome data

**DOI:** 10.1101/2024.09.23.614313

**Authors:** Nicoleta-Denisa Constantin, Adrian Ionascu, Attila Cristian Ratiu

**Affiliations:** Faculty of Biology, University of Bucharest, Intrarea Portocalelor, No. 1-3, 060101, Bucharest, Romania

## Abstract

Rapid evolution of long-read sequencing technologies requires accurate, fast and user-friendly genome assembly and scaffolding tools. In this article we present Digital Scaffolding (dScaff), a bioinformatics tool that performs scaffolding of *de novo* genome assemblies based on reference scaffolds or chromosomes. dScaff makes use of a series of bash and R scripts in order to create a minimal complete scaffold from a genome assembly, with future features to be implemented. Herein, we demonstrate the functionality of dScaff on a novel genome assembly pertaining to a recently sequenced local strain of *Drosophila suzukii*.

## Introduction

The rapid evolution of high-throughput sequencing methods has led to achievable costs for whole genome sequencing (WGS) projects. This present-day reality has stimulated numerous research strategies to make use of WGS in order to analyze various strains of model organisms, as well as specific populations pertaining to species of medical or agronomic interest. In parallel, we are also witnessing computational advances in software development referring to base calling of raw sequencing reads and reference guided or *de novo* reads assembly.

During the recent years, the long-read method developed by Oxford Nanopore Technologies (ONT) became one of the favorite choices for WGS approaches, especially due to its affordability in terms of both infrastructure and reagents costs. In contrast with the classic Next Generation Sequencing (NGS) data comprising short length reads which are not efficient for exposing duplicated or repetitive genomic regions, the use of ONT can bring critical advantages concerning the actual contiguity of a genome assembly.

The proper handling of ONT sequencing protocols can generate substantial amount of data that when exploited by *de novo* assembly heuristics leads to large collections of contigs. Some contigs may contain redundant genomic information or cannot be ascribed to a definite genomic region. Thus, in order to facilitate the analysis/annotation of specific genomic features, one possible strategy is to achieve a minimal assembly made of, if possible, a collection of non-redundant or overlapping contigs assuring a maximum genome coverage.

Repetitive nucleotide sequences that are commonly associated with the centromeres and telomeres heterochromatin, as well as with transposable elements (TEs), are important for genome plasticity and function. Therefore, care should be taken when pursuing annotation endeavors. Moreover, when considerable contig collections result following a given *de novo* assembly that is making use of large ONT data, the probability that the assembly size is impacted by sequence redundancy and/or by the overrepresentation of repetitive elements should to be considered.

With regard to such challenges, we developed the Digital Scaffolding (dScaff) procedure, an original framework for simple and straightforward minimal scaffolding that can even accommodate basic gene annotations of draft *de novo* genomic assemblies. Our approach uses BLAST (Camacho et al., 2009, Boratyn et al., 2013) at its core thus the alignment data that are considered for further analyses are very consistent. Although this procedure is still under development and in this stage may involve some manual steps for output visualization, we demonstrate its use for filtering and ordering specific contigs. Also, we show that dScaff is able to pinpoint specific repetitive genomic regions using nucleotide queries made of reference genome gene sequences (gene queries) available for a fully assembled chromosome.

## Materials and Methods

The efficiency of dScaff was tested on original data gathered for a *Drosophila suzukii* strain derived from a Romanian local population designated ICDPP-ams-1 that was sequenced with ONT and its genome was assembled using Canu software version 2.2 (GenBank/NCBI assembly accession GCA_040114545.1).

For testing, dScaff run a gene queries collection corresponding to gene sequence annotations of chromosome 2L pertaining to the publicly available *D. suzukii* reference genome, Dsuz_RU_1.0 (NCBI RefSeq assembly accession GCF_037355615.1).

The actual scaffolding implementation uses NCBI-BLAST+ version 2.12.0, git version 2.34.1 and seqtk 1.3-r106 Linux packages, R version 4.1.2 and dplyr version 1.1.4, readr version 2.1.5, utils version 4.1.2 and tibble version 3.2.1 R packages. Scaffolding was performed in Linux Mint 21.2 Cinnamon coupled with a i9-13850HX processor, 32 GB DDR4 RAM and 1 TB of storage.

The collection of bash and R (R Core Team, 2023) scripts that makes up dScaff is available on GitHub at https://github.com/DL-UB/dScaff alongside instructions for installing required dependencies and running it.

The bash script that extracts regularly spaced queries from target sequences is available on GitHub at https://github.com/DL-UB/SubSequencesExtractor.

### Premises, implementation and testing of dScaff

Many scaffolding strategies rely on aligning the target contigs with the original collection of reads/contigs or with available reference genomes consisting of fully assembled chromosomes or chromosome specific scaffolds. Consecutively, these procedures use the overlapping regions between contigs that are actually genomic neighbors in order to generate new longer contigs and incidentally decrease/filter their initial arrangement. The amount of working time that has to be considered for such an enterprise can be significant and, in some cases, extensive bioinformatics knowledge is involved.

In its current implementation, dScaff uses BLAST in order to align the draft assembly of interest against the sequences of annotated genes gathered from the corresponding reference genome (if available). When BLAST is employed, if local islands of sequence divergence exist between a long query and the subject sequence the output is not contiguous, but consists in a list of several alignments over the threshold. Hence, by considering from the start that the output will consist of well-defined local alignments generated between a given contig and at least one gene query, dScaff is circumventing the inherent fragmentation of the result in various sub-alignments when there are nucleotide differences between a long query and a comparable or longer subject sequence. Although BLAT (Kent et al., 2002) can stitch disparate sub-alignments, it is prone to make errors when specific sub-sequences are found in several copies within the subject sequence but are uniquely found in the query. Regardless if one uses BLAST or BLAT, if the results are fragmented the specific tabular data that comprise the specific output is hard to interrogate and efficiently exploit in order to recover selected query-subject matches.

The simple and forward dScaff procedure can be functional on any species having at least a draft annotated reference genome available in dedicated public or particular databases. It applies customizable filtering thresholds, retrieves the best query-subject continuous alignments and easily tracks down the collection of alignments between locally arrayed gene queries and a particular tested contig.

We are aware that when considering gene sequences as queries sometimes relevant data cannot be traced, such as some contigs covering chromosomal regions with low gene-density. One of the assumptions of our scaffolding procedure is that the gene density in *D. suzukii* genome is similar to the one particular for *Drosophila melanogaster*. In the latter genome, when considering genomic windows of 50 kb, about 15% of the genes are found in regions harboring less than 5 genes, while the remaining genes are present in regions containing at least 5 genes, up to 29 (Kawahara et al., 2004). The tested *D. suzukii* assembly has an N50 value of approximately 200 kb (data not shown) which leads to the presumption that the corresponding contigs are sufficiently large so that the majority of the genes can be traced to at least one contig. In fact, our selection of contigs facilitated by dScaff mostly comprises contigs harboring more than one gene. Although not detailed here, searching to overcome this apparent limitation brought by region specific gene-density we propose an alternative/complementary strategy so that dScaff is also able to work with collections of discontinuous ordered genomic regions extracted from the reference chromosomes and/or scaffolds (ranked queries). This alternate approach yields a more complex and somehow challenging result to analyze, but provides better coverage of the reference chromosome/scaffold and improves the ability to pinpoint the genomic regions marked by repetitive sequences.

By using the proposed BLAST-based strategy, dScaff can be effectively used on the output provided by other scaffolding strategies such as ntLink (Coombe et al., 2023) or LongStitch (Coombe et al., 2021).

In order to be effective, dScaff requires three major files as input, namely the draft assembly in FASTA format, the specific list or lists of gene sequences from the reference genome assembly and a dataset in TSV format containing the indexes of the respective genes in the considered reference. The latter two files should be downloaded from NCBI database. The dScaff procedure was developed and tested using various draft genome assemblies of a local *D. suzukii* strain, but the results presented herein are based on ICDPP-ams-1 line genome assembly and the publicly available *D. suzukii* reference genome, Dsuz_RU_1.0. If the ranked queries strategy is to be considered, a custom dataset of genomic regions and their corresponding sequences should be used. Additionally, the user has to run a dedicated bash script (SubSequencesExtractor) for taking out the list of ranked queries from a designated target sequence, specifying the length of the query sequences and the distance between them.

Upon running dScaff, the number of the reference chromosomes/scaffolds of the envisioned species will be identified based on the indexed genes table and specific output folders will be created for each chromosome. This step eliminates table entries that are not associated with chromosomes. Next, the gene filtering R script takes as input all the genes for each chromosome and orders these according to the chromosome/scaffold genomic coordinates, regardless of their actual orientation. Therefore, the list will mirror the conventional gene order that is in harmony with the genomic plus strand. The script also contains a variable that can be edited by the user in order to consider only an assortment of genes based on a minimum genomic distance allowed between adjacent genes retained within the resulting filtered list. The intended table is saved in the working folder for each chromosome of the species. Based on this table, the gene sequences are extracted from the command input file using a series of awk, sed and seqtk commands. Next, the draft assembly is hashed for making an NCBI-BLAST+ database (Camacho et al., 2009, Boratyn et al., 2013). The script is optimized to detect the number of available threads on the running machine in order to maximize the performance. Each extracted gene sequence from the previous step is used for running tabular BLAST option (by means of the “-outfmt 6” argument) against the draft assembly database. The results obtained for each individually aligned gene are stored in corresponding .csv files.

Following, the mapping contigs R script takes as input all .csv files and performs strict filtering processes considering two specific parameters: the length of the BLAST alignment and the percent of gene coverage, respectively. The first filtering method retains for further analyses the alignments longer than the standard successive thresholds of gene query coverage and excluded the alignments shorter than 2500 nucleotides. Regarding the percent of gene coverage, our method takes into account successive selection thresholds. Specifically, only a BLAST output file that contains alignments covering more than 50% of the gene length will be further used. If no results fulfill this condition, the threshold is automatically lowered to 40%, and then to a final threshold of 30%. When a particular BLAST output file contains only relatively short alignments (less than 30% of the gene length), the results for that gene are filtered out. Both filtering methods within the mapping contigs R script can be edited by the user, similar to the gene filtering R script.

During the following step, the heuristic pools the filtered BLAST results for each reference genome chromosome. If a certain chromosome from the reference assembly is made of more than one scaffold, dScaff splits the filtered genes table according to each individual scaffold and creates a corresponding output folder. This final output folder is generically named “chromosome” and contains two key CSV files: “indexed_contigs.csv” and “mapped_contigs.csv”. The “indexed_contigs.csv” file has seven columns exhibiting a synthesis of the alignment data produced between contigs and query genes. The first column displays the ordinal number of the contigs specified in the second column and the ranking criteria considers the first occurrence of an alignment between a query gene and a contig. Theoretically, the first contig (1) will be the one that provided a valid alignment with the first gene from the query list. If no contig meets this condition, then the contig 1 will be selected considering a first alignment with the second gene, and so on. When two or more contigs establish a first alignment with the same query gene, they are arbitrary ranked. The third column indicates the number of genes with whom a given contig forms alignments (“genes_hit”). The following columns present the outmost coordinates of the alignment/alignments matching the contig and, respectively, the genomic sequences. For example, when a certain contig forms seven alignments (have seven gene hits) the first contig coordinate will indicate the start of the first alignment with the gene that has the highest or the lowest rank in the query list, depending on the genomic strand that the contig stands for. The last contig coordinate will show the end of the alignment established with the gene query located at the opposite end of the query list in regard with the gene that established the first alignment. Since the order of the genes is not affected by their genomic orientation, the values of the start and end coordinates give insights to the user regarding the genomic strand that is covered by the respective contig. A start coordinate that has a smaller value than the end one will indicate that the contig embodies the genomic plus strand, and alternatively, if the start coordinate has a bigger value, then the contig represents the genomic minus strand. Considering that the genes in the query list are ordered according to the reference strand, the genomic start coordinate will always be smaller than the genomic end coordinate.

The scaffolding procedure is aligning the gene query against the complete contig collection. The eukaryotic genomes are known to harbor many repetitive sequences, especially due to class I and II TEs. *D. suzukii* is no exception and its repeatome represents about 35% of the complete genome (Paris et al., 2020). As a consequence, it is expected that many *D. suzukii* genes are harboring TEs specific sequences, as it is, for example, the case of one of the two *Or22a* gene copies, for which more than half of its sequence is made of two LINE_CR1-4 retrotransposons (according to LBDM_Dsuz_2.1.pri GCF_013340165.1 genome assembly). But LINEs TEs reside in a large variety of genomic regions, within and, especially, outside gene specific sequences, with a tendency towards the heterochromatic regions of the chromosomes. In this context, it is expected that some of the query genes contain repetitive sequences and when such genes generate alignments that pass the dScaff filtering procedure, the respective alignments will be specific for chromosome specific contigs but also for other contigs harboring the same repeated sequence. We remarked that when the “indexed_contigs.csv” file contains sets of contigs sharing the same one or few gene hits this is often the consequence of actually mapping a repeated sequence into multiple contigs with various chromosomal origins. On the other hand, such cases are indicating the chromosomal regions rich in repetitive elements, which could represent valuable information.

In order to offer a schematic visualization of the “indexed_contigs.csv” data, the “mapped_contigs.csv” file presents in a tabular format the specific hits between the contigs (represented on Y axis) and the ordered genes, here designated by their chromosome/scaffold coordinates (represented on X axis). The positive cells indicating a contig-gene correlation are highlighted (Figure 1A). As previously stated, the positive cells are particular to both chromosome specific and non-specific contigs. This simple visual representation can be manually colored in order to emphasize the contigs that will be selected for the intended scaffolding (Figure 1B). At this stage, the user is the one deciding the strategy to follow in order to reach a scaffold. If someone decides that redundant contigs that are located within the same chromosome/scaffold are of interest, the complete mapping representation allows such an approach.

**Figure 1.**
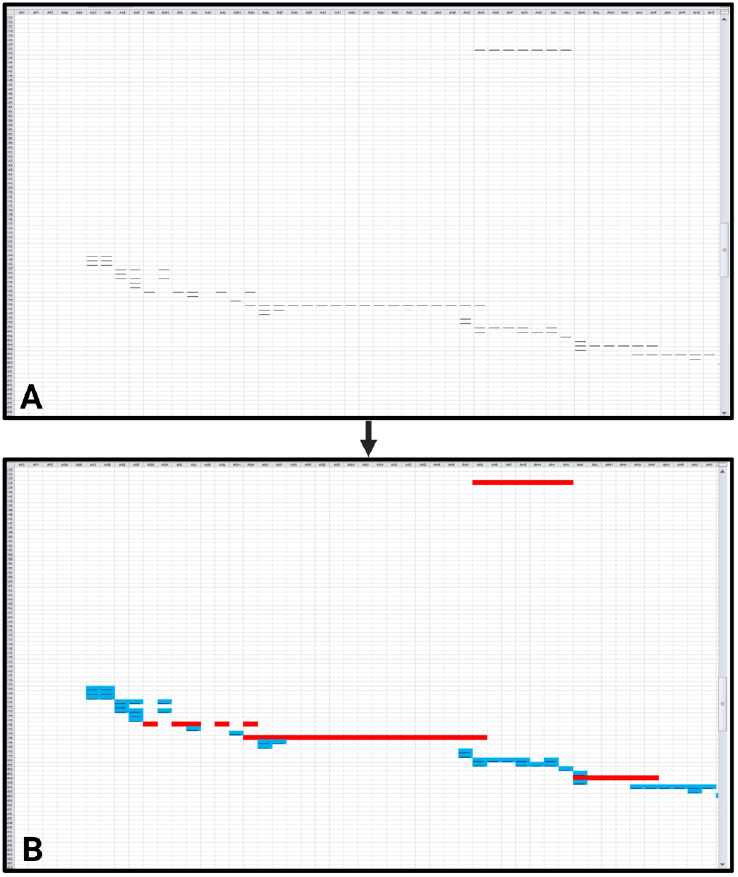
The figure depicts the by default representation of positive blast cells from “mapped_contigs.csv” file (A), which can be manually colored using the Find and Replace function and the Fill/Font Color options of Microsoft Office Excel. A possible choice for highlighting the positive hits that are particular to the contigs forming the minimal scaffold and the remaining ones is to use red and, respectively, blue filled cells (B).

We believe that the most straightforward approach is to attain a so called minimal complete scaffold. We denote with this term the smallest collection of unique contigs that is able to best cover the chromosome/scaffold. In our hands, we chose to mark with red the cells pertaining to the selected contigs and with blue the rest of the positive cells (Figure 1). This strategy leads to an intuitive graphical representation that is efficiently indicating the contig redundancy and the chromosomal regions harboring repeated sequences. When using the gene queries approach it is easy for the user to associate the respective contigs with reference genes. The alternative ranked queries strategy is more suited for analyzing repetitive sequences rich chromosomal regions and requires no gene annotations for the reference assembly. Additionally, it is offering better coverage of the reference chromosomes/scaffolds and longer contiguous BLAST hits.

In the current implementation, dScaff is not able to automatically add colored annotation to the “mapped_contigs.csv” file, therefore manual annotation is required. We recommend converting the CSV file to XLS or XLSX formats before adding any annotations to “mapped_contigs.csv”.

As proof of concept, we primarily tested dScaff performances on the ICDPP-ams-1 genome assembly using the gene queries approach in order to obtain a minimal complete scaffold of chromosome 2L. In Figure 2, the four images are derived from screenshots of the manually annotated “mapped_contigs.xlsx”, where the selected contigs are colored in red and the other positive cells in blue. The regions containing repeated sequences are conveniently indicated by sets of vertically arranged blue cells. Since the right end of 2L chromosome arm is encompassing the centromeric region, it is expected that the signals provided by genomic repeats to be more frequent, which indeed is portrayed in Figure 2D.

**Figure 2.**
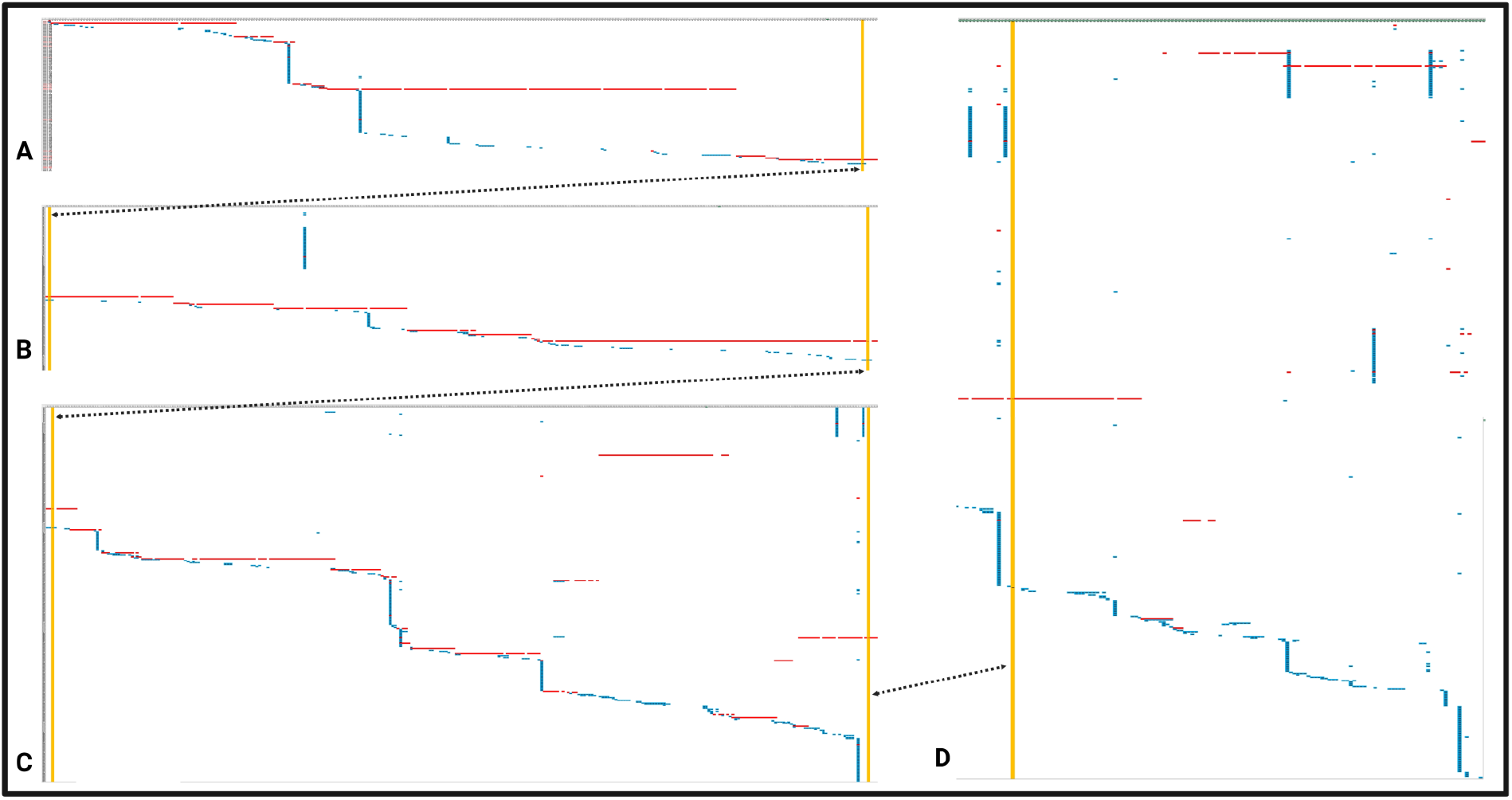
The manually generated minimal scaffold of the 2L chromosome of the ICDPP-ams-1 genotype. The four panels (A to D) are showing the contigs assuring the minimal complete chromosome coverage (the red cells), as well as the other positive cells (blue cells) that represents both redundant and non-specific alignments, the latter probably being a consequence of the repetitive nature of the respective gene queries. The yellow vertical bars are used for establishing a connection between the four panels, the ones residing in the same position being indicated by the double arrowed black dotted line. The many blue cells represented in the D panel are revealing the repetitive nature of the chromosomal sequence neighboring the centromeric region.

Starting from the manually annotated “mapped_contigs.xlsx” established for chromosome 2L of ICDPP-ams-1 genotype, we recovered the complete ordered set of 46 contigs setting up the minimal complete scaffold (Table 1). The smallest contig is the first one, having 28,184 nucleotides, whilst the largest has over 3 million nucleotides. For the first eight contigs we manually analyzed by using BLAST whether the adjacent ones share overlapping sequences and found that except for contig 1, which is completely included in contig 2, all have thousands of nucleotides in common with their genomic assembly neighbor, thus a contiguous region of about 5.8 million nucleotides was exposed. In order to validate the results obtained with the gene queries strategy, we used the same starting data along with the ranked queries strategy. The 4000 nucleotides long query sequences were extracted from the reference sequence of the 2L chromosome with a distance between two adjacent queries of 5000 nucleotides. The manual annotation of the corresponding “mapped_contigs.xlsx” file allowed us to establish a minimal complete scaffold comprising a collection of 51 contigs including 38 contigs selected consecutive to running the gene queries method. The ranked queries approach is not compatible with gene annotations if such data are intended, but was more efficient in identifying longer contigs and pointed out various contigs overlaps characterizing the scaffold generated with our primarily strategy. Moreover, the repetitive nature of specific chromosomal regions is even more evident (Additional file 1).

**Table 1.** The set of contigs pertaining to the minimal complete scaffold generated by means of gene queries strategy. Consecutive to alternatively applying the ranked queries strategy, we discovered that some contigs are actually completely included in larger neighboring contigs (indicated with * symbol) or are actually fully or partially overlapping other contigs, missed by the gene queries strategy (indicated with # symbol).

| Order | Name | Length | Order | Name | Length | Order | Name | Length | Order | Name | Length |
| --- | --- | --- | --- | --- | --- | --- | --- | --- | --- | --- | --- |
| 1* | JBCLUW010002119.1 | 28184 | 13 | JBCLUW010005089.1 | 1006395 | 25# | JBCLUW010006520.1 | 103812 | 37 | JBCLUW010000733.1 | 181089 |
| 2 | JBCLUW010001065.1 | 1711556 | 14 | JBCLUW010001326.1 | 1902342 | 26 | JBCLUW010006519.1 | 512643 | 38 | JBCLUW010002471.1 | 67054 |
| 3 | JBCLUW010001608.1 | 341252 | 15 | JBCLUW010002029.1 | 535342 | 27 | JBCLUW010000783.1 | 123615 | 39 | JBCLUW010001067.1 | 1967922 |
| 4 | JBCLUW010002230.1 | 339634 | 16 | JBCLUW010006025.1 | 460941 | 28* | JBCLUW010000787.1 | 80797 | 40 | JBCLUW010005701.1 | 290679 |
| 5 | JBCLUW010005602.1 | 237944 | 17* | JBCLUW010003327.1 | 30113 | 29 | JBCLUW010001808.1 | 526454 | 41 | JBCLUW010001939.1 | 245168 |
| 6# | JBCLUW010000827.1 | 111384 | 18# | JBCLUW010005619.1 | 53998 | 30 | JBCLUW010001077.1 | 782471 | 42 | JBCLUW010005327.1 | 167602 |
| 7 | JBCLUW010005604.1 | 144826 | 19 | JBCLUW010005620.1 | 3337303 | 31 | JBCLUW010001758.1 | 121533 | 43 | JBCLUW010005520.1 | 493535 |
| 8 | JBCLUW010000160.1 | 3086532 | 20 | JBCLUW010000651.1 | 477662 | 32 | JBCLUW010005574.1 | 323449 | 44 | JBCLUW010004904.1 | 1284471 |
| 9 | JBCLUW010005651.1 | 309004 | 21 | JBCLUW010002223.1 | 349290 | 33* | JBCLUW010002130.1 | 52859 | 45 | JBCLUW010000099.1 | 482170 |
| 10 | JBCLUW010005653.1 | 166449 | 22 | JBCLUW010006181.1 | 62979 | 34 | JBCLUW010001244.1 | 718999 | 46 | JBCLUW010004827.1 | 151105 |
| 11 | JBCLUW010000453.1 | 2128581 | 23 | JBCLUW010000297.1 | 1290666 | 35* | JBCLUW010001748.1 | 65252 |  |  |  |
| 12 | JBCLUW010001453.1 | 220489 | 24 | JBCLUW010000173.1 | 243249 | 36 | JBCLUW010002334.1 | 320211 |  |  |  |

Although revealing the repeated sequences rich genomic regions is of undisputed value, there are situations when only the minimal complete scaffold is of interest. Thus, we are currently developing a prototype additional dScaff R module that automatizes the selection and visual representation of the minimal set of contigs. In short, it uses a heuristic that evaluates the start and stop coordinates of each contig revealed by using gene or ranked queries and selects the longest ones covering a given genomic region. The heuristic removes from the final selection the shortest contig/contigs if several contigs exhibit complete overlap, but retains the partially overlapped adjacent contigs. Although in its current implementation it can be useful for a quick evaluation of the candidate contigs that should form the minimal complete scaffold (Figure 3), it still misses relevant contigs when they generate two or more distanced short alignments with the gene query sequences (data not shown). In summary, the automatic procedure allowed the selection of 38 contigs, 32 being in common with the manually annotated minimal complete scaffold previously described.

**Figure 3.**
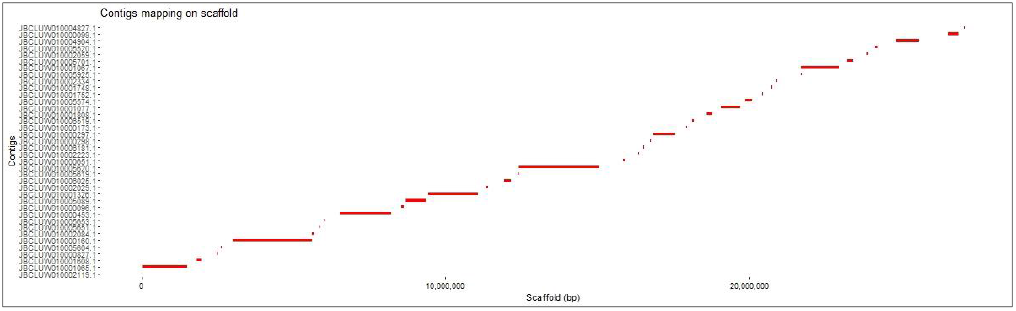
The automatic contigs mapping making a draft variant of a minimal continuous scaffold. The relatively large spaces between adjacent contigs are due to some specific filtration of positive cells based on the alignment length (data not shown).

## Conclusions

The scaffolding strategy that we are propositioning is a straightforward approach that uses a limited number of input files and few computational resources. Also, the procedure requires relatively basic bioinformatics skills as the most stages of our heuristic are automatically completed.

The collection of contigs that could form a minimal complete scaffold is to be manually selected from a very intuitive mapping representation of the positive associations between each contig and the specific queries list. We consider that the manual approach has its strengths, since by differently coloring the desired contigs and the other positive hits, respectively, the user obtains a custom, intuitive graphical representation that successfully highlights the preponderance of repetitive sequences within specific chromosomal regions.

We run original experimental data and sought to expose the strengths and weaknesses of our alternative strategies employing gene or ranked queries, respectively. If best variant of minimal complete scaffold is desired and both gene annotations and precise selection of contigs are sought after the user should complementarily use the two strategies.

In its current variant, dScaff is able to perform a rudimentary variant of automatic graphical representation of the contigs forming the scaffold as well as to extract their identifiers into a corresponding table. Although still in development, we aim to fully integrate this very useful feature in the near future.

As upcoming perspectives, we are planning the implementation of additional features of the dScaff tool, including more sensitive and improved genome scaffolding strategies, publication ready mapping of contigs along the reference chromosomes/scaffolds, exploiting the information contained within the redundant contigs from the draft assembly and annotating genes in the draft assembly based on the refence gene annotations.

## Acknowledgement / Funding

This research was partially supported by the following research project: “Genome sequencing, assembly and annotation of two Romanian populations of *Drosophila suzukii*”, Research Institute of the University of Bucharest, Grant Number: 7419/04.07.2023, Project Director: Attila Cristian Raţiu.

